# A novel 3D atomistic-continuum cancer invasion model: In silico simulations of an in vitro organotypic invasion assay

**DOI:** 10.1101/2020.08.22.263087

**Authors:** Linnea C. Franssen, Nikolaos Sfakianakis, Mark A.J. Chaplain

## Abstract

We develop a three-dimensional genuinely hybrid atomistic-continuum model that describes the invasive growth dynamics of individual cancer cells in tissue. The framework explicitly accounts for phenotypic variation by distinguishing between cancer cells of an epithelial-like and a mesenchymal-like phenotype. It also describes mutations between these cell phenotypes in the form of *epithelial-mesenchymal transition* (EMT) and its reverse process *mesenchymal-epithelial transition* (MET). The model consists of a hybrid system of partial and stochastic differential equations that describe the evolution of epithelial-like and mesenchymal-like cancer cells, respectively, under the consideration of matrix-degrading enzyme concentrations and the extracellular matrix density. With the help of inverse parameter estimation and a sensitivity analysis, this three-dimensional model is then calibrated to an *in vitro* organotypic invasion assay experiment of oral squamous cell carcinoma cells.

## 1 Introduction

The activation of invasion, together with subsequent metastasis, has been identified as one of the hallmarks of cancer (Hanahan and Weinberg, 2000, 2011). Together, these processes account for over 90 % of cancer-related deaths (Hanahan and Weinberg, 2000; Gupta and Massagué, 2006). In epithelial-derived solid tumours, so-called *carcinomas*, the processes of invasion and metastasis involve various phenotypic changes in the initially epithelial-like cancer cells. These are commonly summarised as *epithelial-mesenchymal transition* (EMT) and *mesenchymal-epithelial transition* (MET). Generally speaking, EMT coincides with increased motility through a loss in cell-cell adhesion and a gain in cell-matrix adhesion, as well as increased *matrix-degrading enzyme* (MDE) expression (Micalizzi et al., 2010). Through MET, these processes are reversed. During cancer invasion, complex interactions between cancer cells of various phenotypes and the *extracellular matrix* (ECM) in their tumour-microenvironment typically result in cancer cells moving away from the primary tumour into the surrounding tissue in particular.

Mathematical modelling may provide a complementary approach to help understand the complex mechanisms underlying cancer invasion. However, biologically accurate modelling approaches are crucial to close the often-perceived gap between experimental work and mathematical models. Due to the number of cells involved in cancer invasion, continuum models are a popular and computationally efficient approach to modelling cancer invasion, *cf*. review sections in Franssen et al. (2019); Franssen (2019). This approach can reflect the biology of epithelial-like cancer cells and hence their spatio-temporal evolution well. Moreover, continuous models can be analysed mathematically, see e.g. Ramis-Conde et al. (2008); Andasari et al. (2011); Sfakianakis et al. (2017). However, cancer cells of mesenchymal phenotype play a crucial role in cancer invasion (Godlewski et al., 2010). These cells make up only a small proportion—and hence relatively small number—of cancer cells in the initial tumour (Dongre and Weinberg, 2019). A distinguishing feature of mesenchymal-like cancer cells is their loss of cell-cell adhesion. Hence, it would be biologically inaccurate to represent cells of mesenchymal phenotype through a continuum approach. Modelling cancer cells of various phenotypes along the epithelial-mesenchymal spectrum individually, *cf*. Andasari et al. (2012); Franssen et al. (2019); Franssen and Chaplain (2019), overcomes this problem but the computational cost limits the number of cells that can be modelled. Building on the two-dimensional model by Sfakianakis et al. (2020), we propose a three-dimensional model that represents the spatio-temporal evolution of epithelial-like and mesenchymal-like cancer cells in a biologically appropriate manner while retaining computational efficiency. This is achieved by modelling the epithelial-like cancer cells, which make up the bulk of the tumour, by a macroscopic density profile and their time evolution by a continuum *partial differential equation* (PDE) approach. The more sparsely occurring mesenchymal-like cancer cells through an individual-based *stochastic differential equation* (SDE) approach.

This modelling approach allows us to bridge the often-existent gap between experimental and mathematical work. To demonstrate this, we parametrise the model to accurately represent the invasion of *oral squamous cell carcinoma* (OSCC) cells in an experimental organotypic invasion model proposed by Nurmenniemi et al. (2009). In OSCC, both EMT and MET have been shown to play a crucial role in the local tumour invasion (Chang et al., 2013; Joseph et al., 2018). Through the computational simulations that we carry out, we find that our three-dimensional hybrid atomistic-continuum model of EMT- and MET-dependent cancer cell invasion provides qualitatively and quantitatively biologically realistic outcomes in OSCC invasion.

The remainder of the paper is organised as follows. In Section 2, we explain the role of epithelial- and mesenchymal-like cancer cells, as well as of the transition of cells between those phenotypes, in cancer invasion. Moreover, we describe the biological background, setup and result quantification of the organotypic invasion assay experiments by Nurmenniemi et al. (2009), which we we use to calibrate our model to in the following sections. In Section 3, we introduce the three-dimensional genuinely hybrid model of cancer invasion. Further, we describe how we calibrate it to the particular application of the experiments by Nurmenniemi et al. (2009). In Section 4, we outline the settings, the parameter estimation, and the sensitivity analysis of the simulations of the experiments by Nurmenniemi et al. (2009) that we model. In Section 5, we present and discuss the simulation results. Finally, in Section 6, we discuss the biological implications of our work and planned extensions to the current model.

## 2 Biological background

The invasion of carcinomas into the surrounding tissue is the central biological processed that we model. Due to local constraints e.g. in essential nutrients and oxygen, cancer cells become invasive after the carcinoma reaches a size of approximately 0.1–0.2 cm Folkman (1990). During the invasive growth phase, cancer cells of mesenchymal phenotype are observed in addition to those of epithelial phenotype both *in vivo* (Tsai et al., 2012; Ocaña et al., 2012) and in organotypic assay experiments (Nurmenniemi et al., 2009). In this section, the characteristics of both phenotypes are explained. Moreover, EMT and MET—the processes by which the phenotypes of cancer cells change—are discussed.

Cancer cells adapt to the environmental requirements of their surrounding via changes in phenotype (Jolly et al., 2017). EMT and MET are a canonical group of—at least transiently— observed phenotypic changes that are assumed to be crucial for the spatial spread of cancer (Guo et al., 2012; Ye et al., 2015; Krebs et al., 2017). Various combinations of so-called *EMT-inducing transcription factors* together with a number of extracellular molecules in the tumour microenvironment and related pathways are thought to trigger EMT-/MET-related processes (Jie et al., 2017).

As an outcome of EMT, the cell-cell adhesion between formerly epithelial-like cancer cells, which is predominantly enforced via E-cadherin, gap junctions and tight junctions, is reduced together with their expression of epithelial integrins. Instead, cell-matrix adhesion enhancing molecules like N-cadherin and integrins that are specific to extracellular components on the cell membranes are expressed. Moreover, during EMT the actin cytoskeleton remodels into stress fibres that accumulate at the areas of cell protrusions, and epithelial cytokeratin intermediate filaments are increasingly replaced by vimentin (Micalizzi et al., 2010). As part of this combination of changes, the characteristic polygonal cobblestone-like cell shape of epithelial cells is progressively replaced by a spindle-shaped morphology. Figure 1 schematically shows the changes that cells undergo when switching from an epithelial-like (left) to a mesenchymal-like (right) phenotype. Furthermore, during the EMT process, the motility and invasiveness of the cancer cells are enhanced (Jie et al., 2017; Dongre and Weinberg, 2019) and the cells become increasingly potent at degrading the underlying basement membranes of organs and vessels as well as the ECM via the expression of *metalloproteases* (MMPs) (Dongre and Weinberg, 2019). There are 23 known MMPs (Jackson et al., 2010), which are able to degrade the vast majority of surrounding tissue in humans (Kleiner and Stetler-Stevenson, 1999). These can further be grouped into *soluble* MMPs, like MMP-2 or MMP-9, which are secreted into the surrounding tissue by cancer cells, and *membrane-bound* MMPs, which remain attached to the cell membrane. Amongst the latter, MT1-MMP is particularly well-investigated (Itoh, 2015). Moreover, experimental results by Sabeh et al. (2009) suggest that this membrane-bound MT1-MMP is both necessary and sufficient for cell invasion to occur.

**Figure 1:**
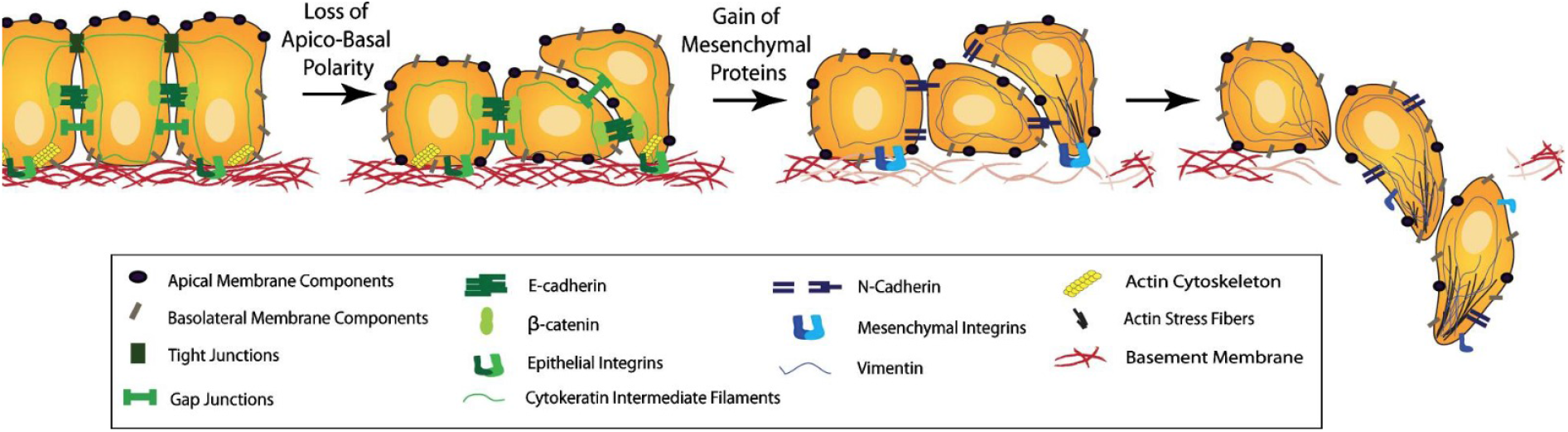
Schematic representation of the EMT (left to right). As an outcome of EMT, the cell-cell adhesion is reduced and invasiveness enhanced through mechanisms explained in the text. Further, cancer cells become more potent at degrading the basement membranes of organs and vessels, as shown towards the right of the figure, as well as the ECM in general. This allows the mesenchymal-like cancer cells to invade the surrounding stroma. During MET, which can be understood by reading the figure right to left, these changes in phenotype are reversed. Reproduced from Micalizzi et al. (2010) with permission from Springer. ((pending))

Through MET the phenotypic changes induced by EMT can be reversed. Thus—generally speaking—MET causes the cells to become less motile and invasive while enhancing their proliferative potential.

A non-reversible, stable transition from an epithelial to a mesenchymal phenotypic state, which had formerly been assumed to be the only possible outcome of EMT, has recently been shown to actually be rare during carcinogenesis (Dongre and Weinberg, 2019). Instead, a plasticity to switch between the two phenotypic states through EMT and MET is suggested to exist for most invading cancer cells (Thiery, 2002; Tsai and Yang, 2013).

During local cancer invasion, some cancer cells in the tissue have been found to be of mesenchymal phenotype (Friedl and Wolf, 2003). Hence, EMT in at least a subset of the initially epithelial-like cancer cells at the primary site is a prerequisite for invasion (Francart et al., 2018; Pastushenko and Blanpain, 2018). Migrating cells usually employ their acquired mesenchymal traits, i.e. the decrease or loss in cell-cell adhesion and increase in cell-ECM adhesion and in MDE-expression, to invade (Friedl and Wolf, 2003; Bill and Christofori, 2015). The proliferation-enabling MET, on the other hand, is involved in metastatic colonisation in most carcinomas. In fact, it has been suggested that stable mesenchymal-like phenotypes without any MET potential cannot succeed in metastatic re-seeding (Ocaña et al., 2012; Ruscetti et al., 2015; Kröger et al., 2019).

The focus of this paper lies on the invasion of OSCC, the most common type of *head and neck squamous cell carcinoma* (HNSCC) (Kim et al., 2019). In HNSCC, distal organ metastasis occurs rarely compared to other cancers. Instead, local progression is a major cause of HNSCC-related mortality (Chang et al., 2013). A study by Chang et al. (2013) suggests that the mechanism behind the aggressive local spread observed in HNSCC is the induction of MET in the tumour microenvironmemt rather than at the distal sites, as typically observed in carcinomas (Dongre and Weinberg, 2019). This results in the inhibition of migration for HNSCC cells— such as the HSC-3 cells modelled in this paper—through connective tissue growth factors in the microenvironment of a primary tumour. Thus, MET enables faster growth of the formerly individually occurring cancer cells that were previously of an invasive mesenchymal phenotype. An often-observed phenomenon in OSCC, as well as other types of carcinomas (Japanese Gastric Cancer Association, 2011; Ito et al., 2012; Masuda et al., 2017), is the occurrence of “islands” of cancer cells outside of the main body of the tumour (Nurmenniemi et al., 2009; Almangush et al., 2018). Notably, in OSCC, the “island” count has, through a number of studies, been confirmed to be a reliable and simple prognostic marker that correlates with poor prognosis (Almangush et al., 2018).

### 2.1 An *in vitro* organotypic invasion assay study

In this section, we introduce the three-dimensional organotypic invasion assay experiment by Nurmenniemi et al. (2009). This is the experiment that we subsequently reproduce through the mathematical model for cancer invasion of the ECM, showing that this model provides biologically accurate predictive results. The experiment by Nurmenniemi et al. (2009) studies the invasion of human tongue squamous cell carcinoma cells of the cell line HSC-3 into uterine leiomyoma tissue. As they show in the corresponding paper, this mimics the *in vivo* invasion of the tumour microenvironment in OSCC.

In what follows, a description of the experimental setup of the invasion organotypic assays in Nurmenniemi et al. (2009), their experimental results and the methods of result quantification is given.

For the organotypic culture, Nurmenniemi et al. (2009) selected only non-degraded human uterine leiomyoma tissue, in the preparation of which any areas with macroscopically heterogeneous tissue were omitted. The suitable tissue was cut into 3 mm thick slices. From these slides, discs of 8 mm diameter were punched. Then, 7 × 10^5^ human tongue squamous cell carcinoma cells of line HSC-3 were allowed to attach to the top of each myoma disc overnight. Subsequently, the myoma discs were transferred onto uncoated nylon discs that rested on curved steel grids in 12-well plates with sufficient volume of media.

At days 2, 8 and 14, the organotypic cultures, which all stemmed from the same myoma to minimise differences in tissue, were formalin-fixed. Then they were dehydrated, bisected and embedded in paraffin. Next, they were sectioned into slices of 6 µm thickness and immunostained according to the question the authors sought to address. For the main invasion experiment, which our model focusses on, pancytokeratin AE1/AE3 was used, which stains epithelial-derived HSC-3 cells brown.

#### Quantification of experimental results

The top row in Figure 6 shows microscopic fields with pancytokeratin AE1-/AE3-stained organotypic assays, that initially consisted of a single HSC-3 cell layer on top of visually homogeneous myoma tissue, at day 2, 8 and 14 (left to right).

Quantitatively, invasion results were measured using three “norms”: the *maximal invasion depth*, the *invasion area* and the *invasion index*, as shown in Figure 2 (left to right). To determine the maximal invasion depth for each slice, the distances of the three HSC-3 cells that had invaded furthest from the myoma surface—measured perpendicularly to the top edge of the microscopic field—were measured using Fiji software, as shown in Figure 3, and the mean of the distances was calculated, as described in Åström et al. (2018). This was repeated for 3 to 8 slices from the same myoma disc and then averaged. Using this method in this experiment, the maximal invasion depth was found to be 547 µm, with an interquartile range of 61 µm. To calculate the invasion index, *cf*. Nyström et al. (2005), Nurmenniemi et al. (2009) first quantified the area of the upper non-invading cell layer, which corresponds to the respective area coloured in white in Figure 4 in each microscopic field, as well as the area occupied by the sum of the remaining invading epithelial-like HSC-3 cells, which is highlighted in red in the same figure. These measurements were, again, taken from 3 to 8 slices of the same myoma to determine the mean area of the upper non-invading cell layer 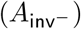 and the mean invading cell area 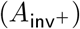, respectively. The invasion index (*I*) was then calculated as

**Table 1:**
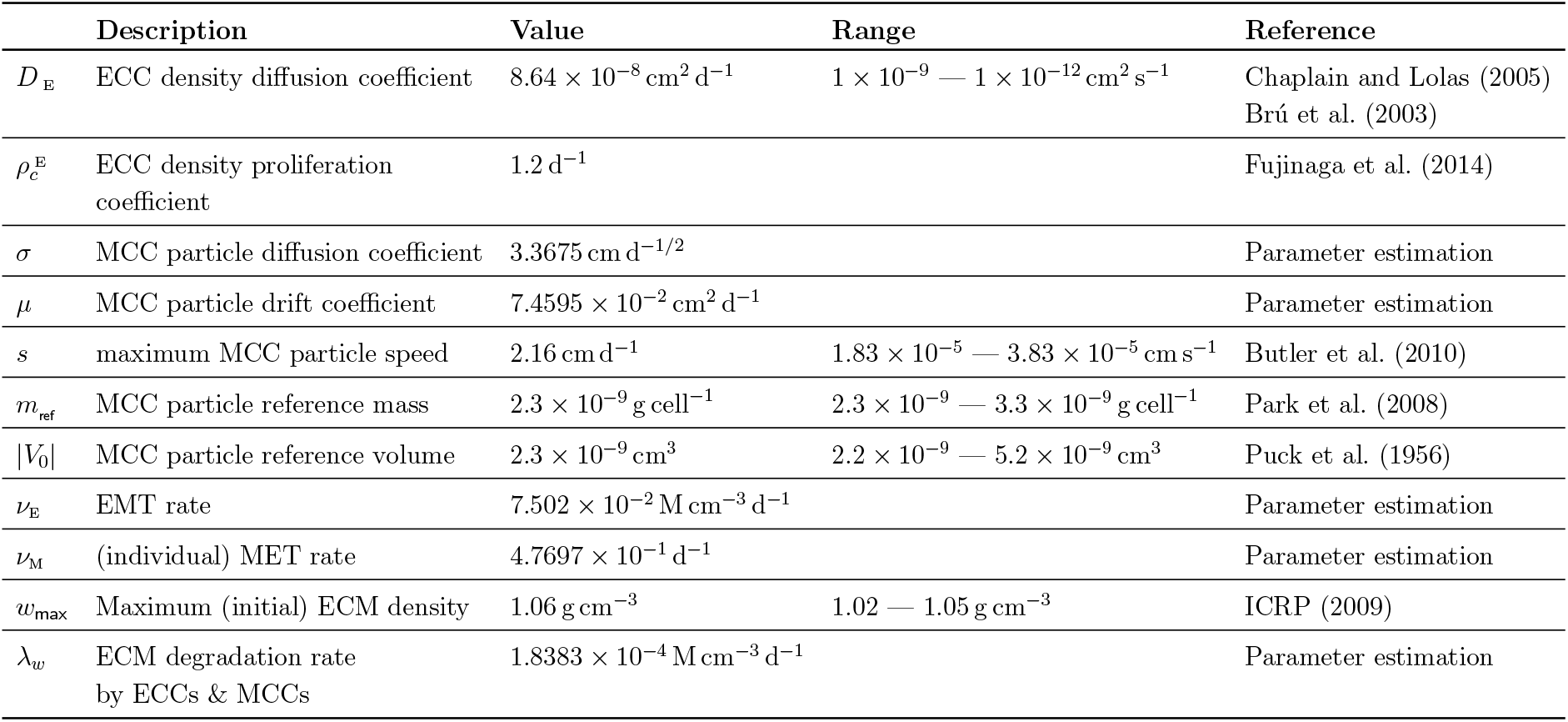
Parameter settings for the simulations. Epithelial-like HSC-3 cells and mesenchymal-like HSC-3 cells are abbreviated *ECC* and *MCC*, respectively.

**Figure 2:**
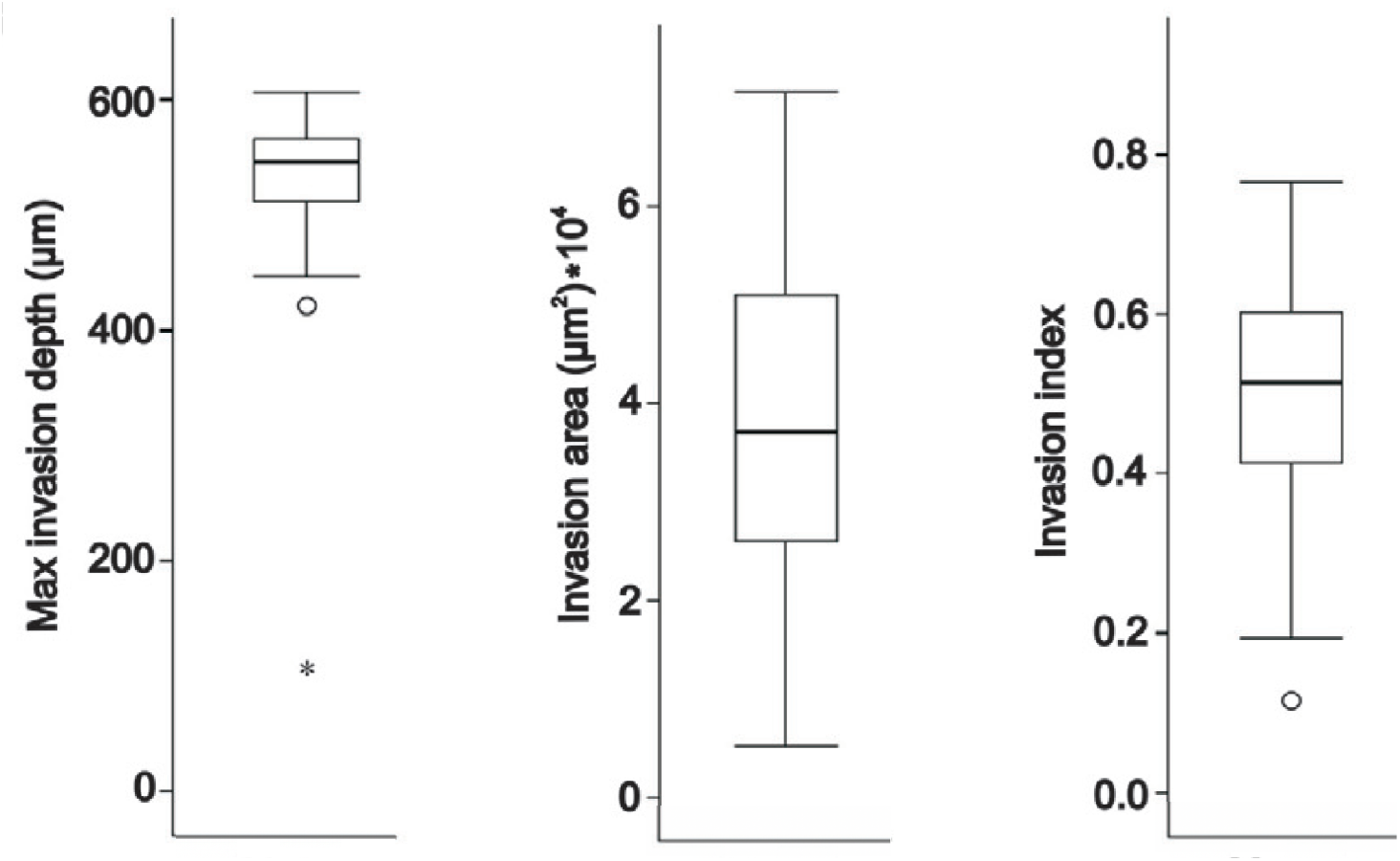
Quantification of the organotypic invasion in Nurmenniemi et al. (2009) and in Section 2. **1**. The measurements of maximal invasion depth, invasion area, and invasion index were taken from myoma tissue on day 14 of the invasion. The central rectangles span from the first to the third quartile. The segment inside the rectangle shows the median. The whiskers represent the locations of the respective minimum and maximum. Suspected outliers are indicated by a circle and outliers by a star. The results for the maximal invasion depth consist of at least three measurements, cf. Figure 3, of two to eight slices from two to four independent assays. For the invasion area and index, one measurement per representative area was taken from each of the two to eight slices from the two to four independent assays. This figure is modified from Nurmenniemi et al. (2009) with permission from Elsevier.((pending))

**Figure 3:**
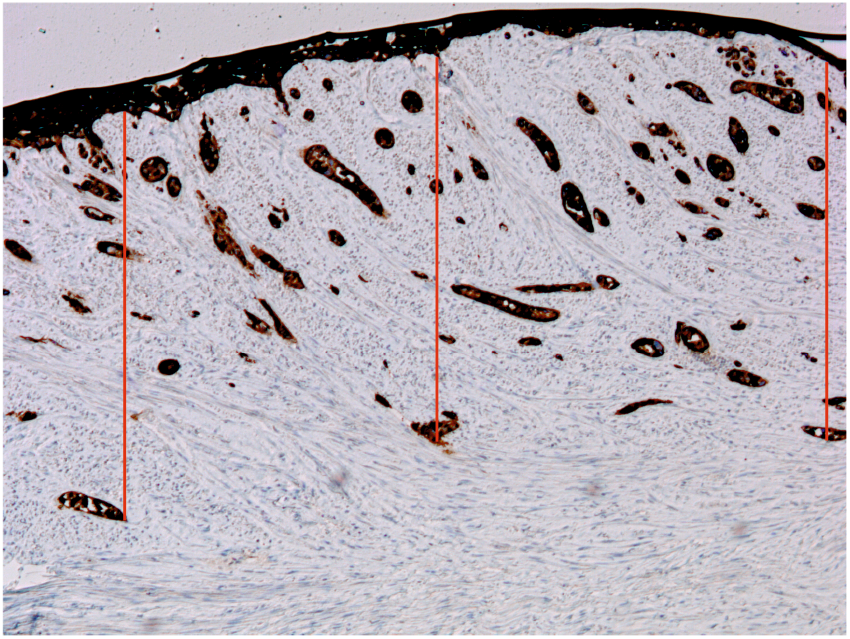
Maximal invasion depth. The invasion distances of the three epithelial-like HSC-3 cells that invaded furthest into the myoma—measured perpendicularly to the top edge of the microscopic field—were measured as indicated by the red line. Next, their mean was calculated as described in Åström et al. (2018).

**Figure 4:**
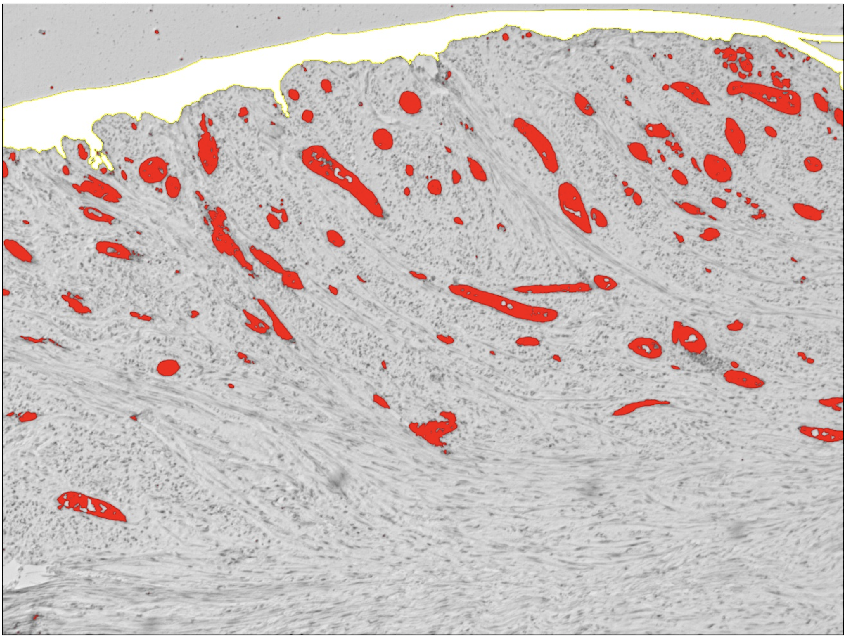
Invading vs. non-invading cell area. The area of the upper non-invading epithelial-like HSC-cell layer is shown in white; the area that is occupied by invading epithelial-like HSC-3 cells is shown in red. Cells of mesenchymal-like phenotype were not accounted for in the determination of the respective areas.

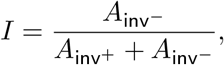

which gave *I* = 0.51 [0.41, 0.60] in this particular experiment.

## 3 Model description

The model we propose is a three-dimensional, hybrid atomistic-continuum that accounts for the collective epithelial and the individual mesenchymal invasion strategies, as well as for the EMT and MET between these two phenotypic cell states. The proposed model is based on a two-dimensional cancer invasion model introduced in Sfakianakis et al. (2020). Besides the extension to a third dimension, the current model features the production and action of membrane-bound MT1-MMPs, the migration of the mesenchymal-like cancer cells and, most importantly, the transition between the two phenotypic states. Moreover the parameters of the model are calibrated via parameter estimation and sensitivity analysis to accurately represent the organotypic HSC-3 invasion assay experiments by Nurmenniemi et al. (2009).

We formulate a hybrid atomistic-continuum model in the sense that the epithelial-like cancer cells are described by a continuum density distribution whereas the mesenchymal-like cancer cells are modelled as a collection of isolated cells. To emphasize their discrete nature, we from now on refer to the latter as *cell-particles*.

Epithelial-like cancer cells typically appear as cell sheets with strong intercellular adhesions and, accordingly, their time evolution is modelled by a system of PDEs. The cell-cell adhesions are diminished through EMT. This causes resulting mesenchymal-like cancer cells to typically appear individually, *cf*. Figure 1. Their time evolution is modelled by a system of SDEs. The cell-cell adhesions and hence the epithelial phenotypic state can be regained via MET. These PDE and SDE submodels are coupled through the *density-to-cell-particle* and *cell-particle-to-density* operators, *cf*. Appendix A, that model the transitions from one phenotypic state to the other.

### Density-based submodel

Through the density description and a system of macroscopic deterministic PDEs, in particular, we model the spatio-temporal evolution of the epithelial-like cancer cells, the membrane-bound MT1-MMPs and the ECM. We assume that the epithelial-like cancer cells compete for space and resources with the mesenchymal-like cancer cells and the ECM. No active migration of the epithelial-like cancer cells is assumed—neither in the form of a directed response to extracellular chemo- or haptotaxis cues nor in the form of random cell migration. Still, we make the assumption that the epithelial-like cancer cells proliferate and that this introduces small mechanical pushing forces between them. This is incorporated into the model through a small diffusion term.

We denote by Ω ⊂ ℝ^3^ a Lipschitz domain suitable for the experimental settings and by *c*_E_(**x**, *t*), *c*_M_(**x**, *t*), *m*(**x**, *t*) and *w*(**x**, *t*), where **x** ∈ Ω and *t* ≥ 0, the densities of the epithelial-like cancer cells, the mesenchymal-like cancer cells (whenever applicable), the non-diffusible MT1-MMPs and the ECM, respectively. The subscript E indicates the epithelial phenotype and M the mesenchymal phenotype throughout the paper.

The mesenchymal-like cancer cells are primarily described through their cell-particle formulation. However, the mesenchymal-like cancer cells participate in the time evolution of the epithelial-like cancer cells, which are described by a density formulation, via their density formulation *c*_M_. This is obtained by the *cell-particle-to-density* process explained in Appendix A.

The above considerations are incorporated in the following PDE that governs the spatio-temporal evolution of the density *c*_E_ of the epithelial-like cancer cells:

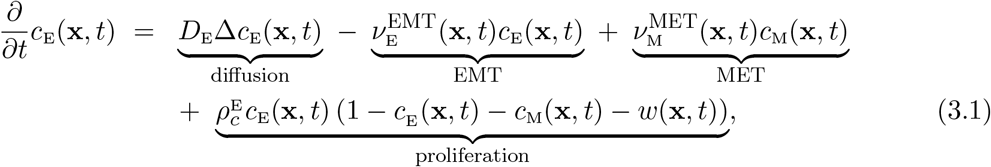

where 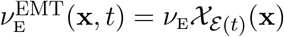 and 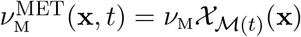, with *ε* (*t*), *ℳ*(*t*) ⊂ Ω, and 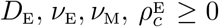. For simplification, we assume that the EMT takes place in randomly chosen sets, denoted by *ε* (*t*) ⊂ Ω. We understand *ε* (*t*) as the set union of a number of sub-sets each having the size of one biological cell, *cf*. Appendix A. Similarly, we make the simplifying assumption that the mesenchymal-like cancer cells, modelled as isolated cell-particles, undergo MET in a random fashion. This gives rise to *ℳ*(*t*), which is another union of sub-sets, each of the size of a single cancer cell, *cf*. Sfakianakis et al. (2020).

We also assume that the mesenchymal-like cancer cells produce non-diffusible MT1-MMPs that are expressed on the cell membranes. Furthermore, for simplicity, we let all mesenchymal-like cells express the same fixed number of MMPs on their membrane. We denote by *m* the density of MMPs and account for them through

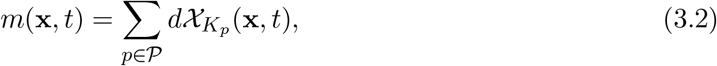

where *K*_*p*_ represents the physical space occupied by the mesenchymal-like cells with index *p*; 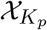 is the corresponding characteristic function; and *d* represents the “number” of MMPs expressed on the cells’ membrane.

The ECM is assumed to be an immovable component of the system that neither diffuses nor otherwise translocates. It is moreover assumed not to be reconstructed in any way. It is degraded by the action of the mesenchymal-like cell/MMP-complexes. Altogether the ECM is described by a (non-uniform) density profile that evolves in time according to

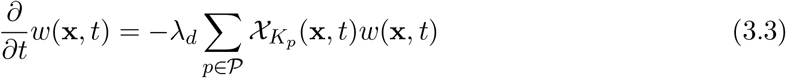

where *λ*_*d*_ = *λ*_*w*_ *d* is the effective degradation rate of the ECM.

### Cell-particle based submodel

The spatio-temporal evolution of the mesenchymal-like cancer cells is dictated by the cell-particle based submodel. Similarly to the rest of the model, the methods and techniques used here are motivated by the work in Sfakianakis et al. (2020) and the references therein.

The mesenchymal-like cancer cells are modelled as a collection of isolated cell-particles that migrate through the tissue while performing a *biased random motion*. The biased part of their motion is due to their haptotactic response to gradients of the ECM, while the random part is understood as a Brownian motion. In the current model no interaction between the mesenchymal-like cells is assumed.

At any given time *t*, we consider a system of *N* = *N* (*t*) ∈ ℕ mesenchymal-like cancer cells, indexed by *p* ∈ *P* = {1, …, *N*}, and we account for their positions **x**_*p*_(*t*) ∈ ℝ^3^, and masses *m*_*p*_(*t*) ≥ 0. Their migration is modelled by a system of SDEs—one SDE for each cell-particle— and is comprised of a combination of two independent processes: a directed motion component that represents the haptotactic response of the cells to gradients of ECM-bound adhesion sites, and a random motion component that is modelled as a Brownian motion:

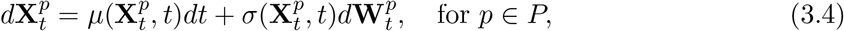

where 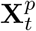 represents the position of the cell-particles in physical space (here ℝ^3^), and 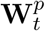 is a Wiener process. The *drift* term *µ* encodes the directed (or biased) part of the motion, whereas the *diffusion* term *σ* encodes its random component.

The mesenchymal-like cancer cells participate in several dynamical processes, e.g. in the EMT and MET, in the proliferation of the epithelial-like cancer cells, in the production of MMPs, and in the degradation of the ECM. Yet, the cell-particle migration equation (3.4) does not include any reaction terms. Instead, these are accounted for in the following way: as mesenchymal-like cancer cells undergo MET and acquire epithelial-like character, they are transformed to a density profile via the *density-to-particle* operator, *cf*. (3.7) and Appendix A. This additional epithelial-like cell density augments the existing epithelial-like cell density and participates in the dynamics modelled through the system of equations (3.1)–(3.3). Conversely, parts of the epithelial-like cancer cell density undergo EMT and transform into mesenchymal-like cancer cells, which are initially described by a density profile. This density is then transformed into particles via the *particle-to-density* operator defined in (3.7) and Apendix A.

### Hybrid formulation of cancer cells

We assume that the domain Ω is sufficiently large and regular to be uniformly partitioned as Ω = ∪_*i*∈*I*_ *M*_*i*_, where every *M*_*i*_, *i* ∈ *I* is a translation of a generic cube *K*_0_ ⊂ ℝ^3^, representing the volume occupied by a single biological cell. This partition allows to represent every scalar (measurable) density function *c* : Ω × (0, ∞) → ℝ by its simple-function decomposition

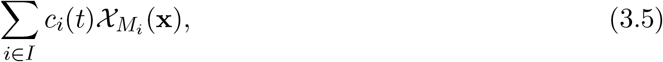

where 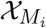 is the characteristic function of *M*_*i*_ ⊂ Ω, and *c*_*i*_(*t*) the mean value of *c*(·, *t*) over *M*_*i*_, i.e.

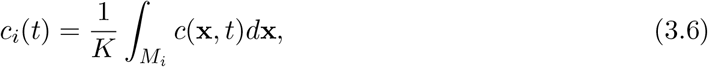

where *K* is the volume of *K*_0_ and, effectively, of *M*_*i*_.

Accordingly, the hybrid description upon which this work is based reads

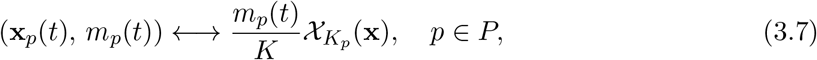

where **x**_*p*_(*t*) and *m*_*p*_(*t*) represent the time-dependent position and mass of the cell-particle; *K*_*p*_ is the translation of the generic cube *K*_0_ with centre **x**_*p*_; and *K* = |*K*_*p*_| = |*K*_0_| is the volume of these cubes. Based on (3.7), the transition between the two cell phases is conducted by the *particle-to-density* and the *density-to-particle* transition operators, see Appendix A.

## 4 Model initiation and parametrisation

We set the state variables and parameters of our model according to the experimental settings introduced for the invasion assay experiments in Nurmenniemi et al. (2009). In what follows, we explain how we reproduce the experiments in detail. Note that all model simulations, including the parameter estimation and the sensitivity analysis, were conducted using MATLAB (2019), and all visualisations with ParaView (Ahrens et al., 2005).

### 4.1 Initial and boundary conditions

The ECM is represented by a three-dimensional landscape that is initially randomly structured with values between a biologically relevant minimum and maximum ECM density, in accordance with ICRP (2009). The construction of the initial ECM density is computational, inductive, and based on discrete principles. Namely, an initial 8 × 8 × 8 random matrix with normally distributed values between the predefined minimum and maximum density values is refined through bisection to 16×16×16 then to 32×32×32 and so on until the computational resolution of the domain Ω is reached. At every refinement stage, the new values are interpolated from the previous ones with the addition of some Gaussian noise. A two-dimensional representation of this procedure is shown in Figure 5.

**Figure 5:**
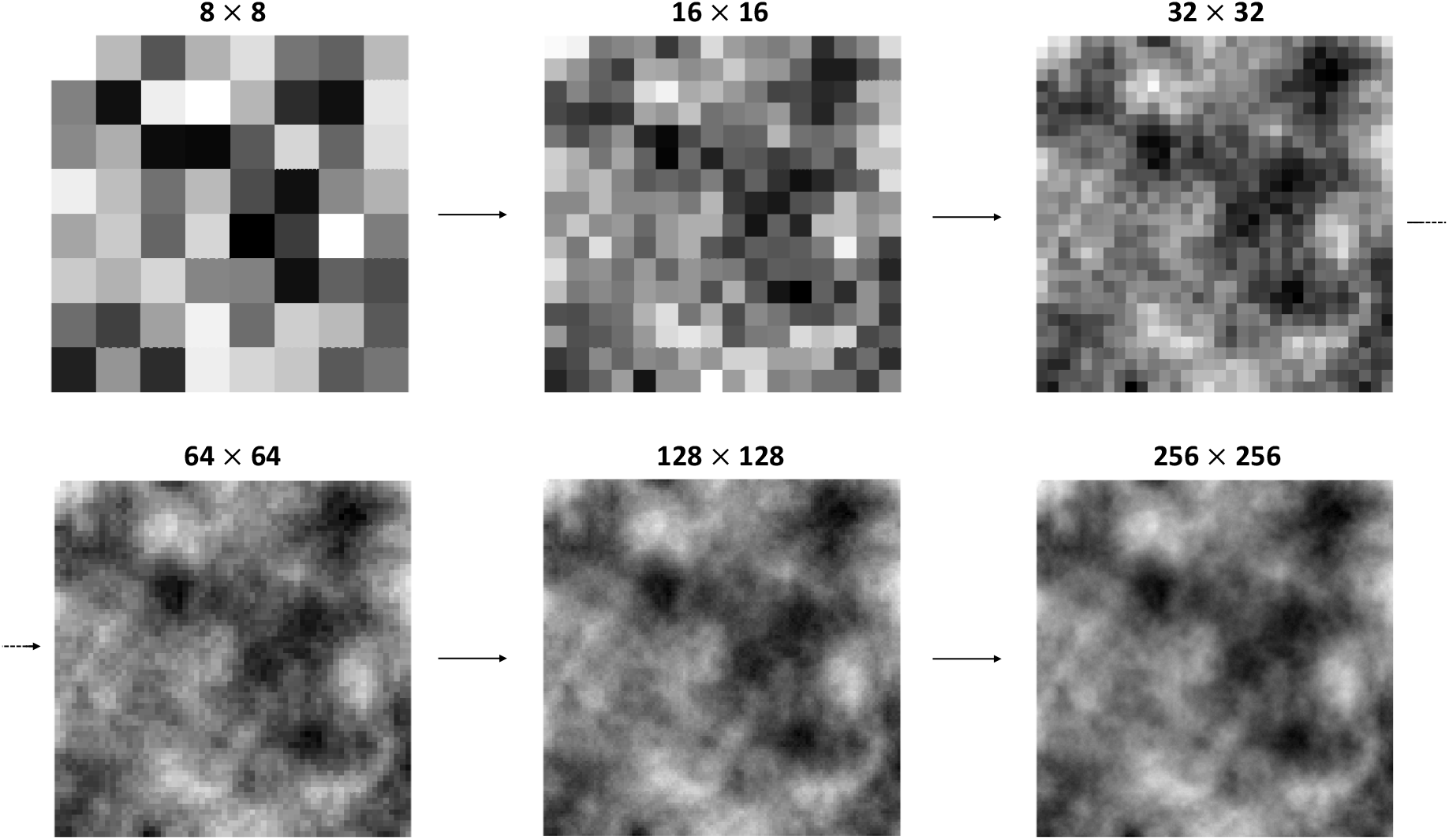
Construction of a sample initial ECM density distribution in two dimensions. An initial 8 × 8 random matrix (top left) is refined progressively to a 256 × 256 matrix (bottom right). At every refinement step, the density values of the refined matrix are obtained from the predecessing coarser matrix by interpolation and periodic extrapolations. The result is a sample two-dimensional initial ECM density distribution with values between w_min_ and w_max_, cf. Table 1 and (ICRP, 2009). In the current work, the corresponding process was carried out in three dimensions with a final refinement of 64 × 64 × 64.

In accordance with the experiments by Nurmenniemi et al. (2009), where a layer of 7 × 10^5^ epithelial-like cancer cells and no mesenchymal-like cancer cells were placed on top of the myoma discs, we consider a single layer of epithelial-like cancer cells placed onto the upper surface of the ECM.

Throughout this work, we consider zero Neumann boundary conditions for the epithelial-like cancer cell density. No boundary conditions are assumed for the ECM as it has been modelled as an immovable component of the system. We do not impose any boundary conditions on the MMPs since the particular family we consider in this model, MT1-MMP, is bound to the cancer cell membrane. Furthermore, no mesenchymal-like cell-particle is allowed to leave the domain Ω. Instead, every time a cell-particle escapes, it is returned to its last known position within Ω and is allowed to resume its biased random motion (3.4).

### 4.2 Model parameters

To ensure that our simulations are biologically realistic, we use parameter values from the literature wherever possible. These are summarised in Table 1 together with the corresponding literature sources. Still, five of the parameters could not be obtained from the literature and had to be indirectly inferred. To this end, and due to the inherent stochasticity of the model, we used a combination of global and local optimisation techniques, which we augmented with a sensitivity analysis.

#### 4.2.1 Parameter estimation

The five parameters that need to be indirectly inferred are the ECM degradation rate *λ*_*w*_, the drift and diffusion coefficients of the mesenchymal-like cancer cell-particles (*σ* and *µ*), and the EMT and MET rates *ν*_E_ and *ν*_M_, *cf*. Table 1.

As the global optimisation method we use *enhanced scatter search* (eSS) (Egea et al., 2009). It belongs to the wider class of stochastic global optimisation methods called *metaheuristics* (Glover and Kochenberger, 2006). Like other stochastic optimisation methods, eSS draws an initial diverse population of guesses out of the parameter space and conditionally initiates intense local searches. For the local optimisation we implement the *interior point method, cf*. Byrd et al. (2000); Karmarkar (1984). This is an iterative linear and non-linear convex optimization method that achieves its goal by going through the middle of the bounded multidimensional polyhedron in the parameter space. Due to the robustness of the method, it is well-suited for a problem of mixed stochastic-deterministic nature like the one addressed in this paper. For the metaheuristic part, we use the *metaheuristics for bioinformatics global optimization* (MEIGO) toolbox (Egea et al., 2010).

The set of five parameters to be inferred is denoted as

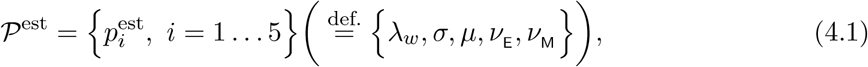

and is estimated through the minimisation of the discrepancy/error between the experimental measurements and the model predictions. This error is measured by the *objective functional*

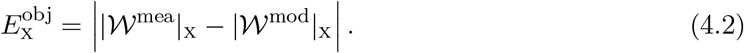

The “norm” |*W* ^mea^|_X_ denotes the experimentally measured quantities, indicated here by the subscript | *·* |_X_. |𝒲^mod^|_X_ on the other hand denotes the corresponding quantity predicted by the model. For the purpose of this work, we considered “norms” such as the *maximal invasion depth* of the epithelial-like cancer cells, their *invading cell area*, and combinations of these, *cf*. Section 5.1, in accordance with Nurmenniemi et al. (2009). The experimental meaning of these quantities is explained in Section 2.1, and in Figures 3 and 4.

#### 4.2.2 Sensitivity analysis

Each of the model parameters, especially the ones inferred by the parameter estimation, has a different impact on the dynamics of the model. A profound understanding of the effect of the model parameters is crucial both for drawing biological conclusions and for quantifying the corresponding biological processes. Moreover, the sensitivity analysis is useful for the calibration and further development of the model.

We study the qualitative and quantitative impact of the parameters by performing a *local sensitivity analysis*. We first consider a particular experimental setting and a reference parameter set. This parameter set is obtained through the previously discussed parameter estimation methods. We then vary the parameters in the reference set one after the other and compute the corresponding numerical solutions of the model. As the problem is stochastic in nature, we repeat every numerical experiment (i.e. with the same settings and parameters) several times and average the corresponding results. We compare each of these solutions to the reference solution through the chosen objective functional (4.2). This way, we quantify the effect that the variation of this parameter has on the solution.

In more detail, we denote the parameters whose effect we study as

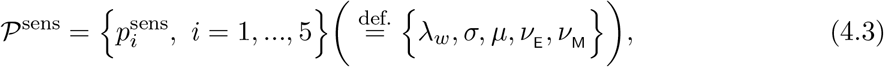

and the *reference parameter set* as

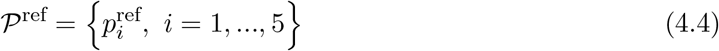

and use it to compute the reference solution 𝒲^ref^. *Ceteris paribus*, we perturb the reference parameters 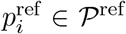, which gives new values 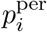. For each perturbation of the parameters, the new parameter set is denoted as

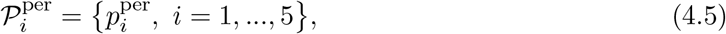

which differs from the reference set 𝒫^ref^ (4.4) only for the parameter *i*. With the perturbed parameter set 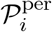, and otherwise the same model conditions (initial, boundary, etc.), we compute the corresponding solution 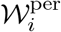. We then compare 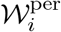 to the reference solution 𝒲^ref^. For this, we use the *sensitivity function*

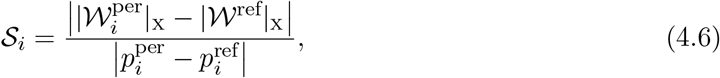

where | · |_X_ denotes a suitably chosen “norm”, *cf*. Section 5.1. In essence, as defined in (4.6), 𝒮_*i*_ represents the absolute rate of change of the solution 𝒲, in the sense of the “norm” | · |_X_, with respect to the parameter *i* around its reference value 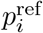.

The *local sensitivity* follows from accounting for all the parameters of 𝒫^sens^. It can be used to deduce qualitative and quantitative biological information. For instance, it lets us rank the influence of various parameters on the system. This, in turn, enables us to gain a more comprehensive understanding of how the different parameters and their variation impact the simulation results.

## 5 Simulations and results

To perform the numerical simulations, we replicated the experimental setting used in Nurmenniemi et al. (2009) as closely as possible. Accordingly, we reconstructed qualitative features of the myoma organotypic assays described therein by using results by ICRP (2009) on the average density of human uterine ECM and by reproducing the homogeneity of the ECM. This is described in Section 4 and Figure 5. Secondly, we have conducted our simulations over a cubic domain of size 8500 µm × 8500 µm × 8500 µm, similar to the assays used in Nurmenniemi et al. (2009). In the experiments, this domain was cylindrical with a diameter of 8000 µm and a height of 3000 µm. However, the 6 µm thick slices that were ultimately analysed in Nurmenniemi et al. (2009) were only of size 600 µm × 600 µm and were taken perpendicular to the myoma disc surface. As Figure 6 shows, we examined slices of dimension 600 µm × 600 µm × 6 µm from the larger three-dimensional domain. The initial conditions we considered correspond to those in the experiments by Nurmenniemi et al. (2009) for which 7 × 10^5^ epithelial-like and no mesenchymal-like cancer cells were placed on top of a myoma disc as a single layer.

Primarily due to the diffusion term, (3.1) causes a slight propagation of the epithelial-like cancer cell front in the mathematical model. This becomes apparent mostly in the earlier stages of the time evolution before new cancer-cell “islands” are formed. At the same time, isolated mesenchymal-like cancer cells arise due to EMT. These are modelled as cell particles whose migration is dictated by (3.4) and are indicated by red dots in Figure 7.

**Figure 6:**
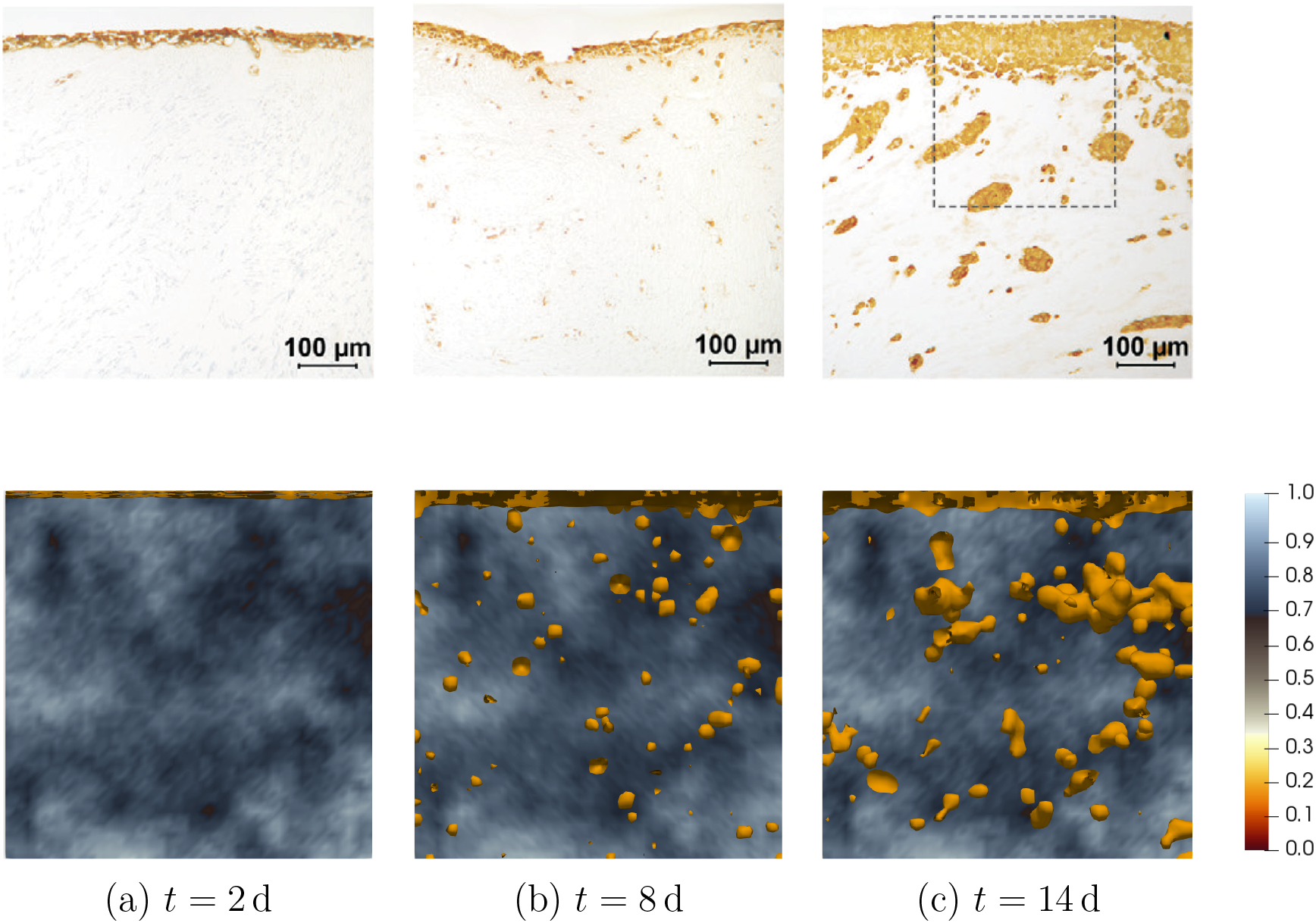
Experimental in vitro versus model simulation results for HSC-3 myoma invasion. The spatio-temporal evolution of an initially uniformly dense epithelial-like cancer cell population placed on top of an ECM of heterogeneous density is depicted after 2, 8, and 14 days. Experimental results of HSC-3 myoma invasion assays by Nurmenniemi et al. (2009) are shown in the top row of panels and the corresponding sample simulation results in the bottom row. All panels show slices of a three-dimensional assay; each of size 600 µm × 600 µm and 6 µm of thickness. In the simulation results (second row), the epithelial-like cancer cells density is represented via the yellow-orange isosurfaces corresponding to 0.1 (and higher) of the average tumour density. The colour bar corresponds only to the density of the ECM; the maximum value being the biological relevant 1.06 g cm^−3^, cf. Table 1. EMT spawns mesenchymal-like cancer cells (not depicted here), which escape the main body of the tumour and invade the ECM more rapidly than the slowly diffusing epithelial-like cancer cells. The reverse process, MET, gives rise to the epithelial-like cancer cell “islands” observed in the middle and right panels. As seen in the 14 d panels, these “islands” eventually reconnect with the main body of the tumour. The top row panels are modified from Nurmenniemi et al. (2009) with the publisher’s permission ((pending)).

**Figure 7:**
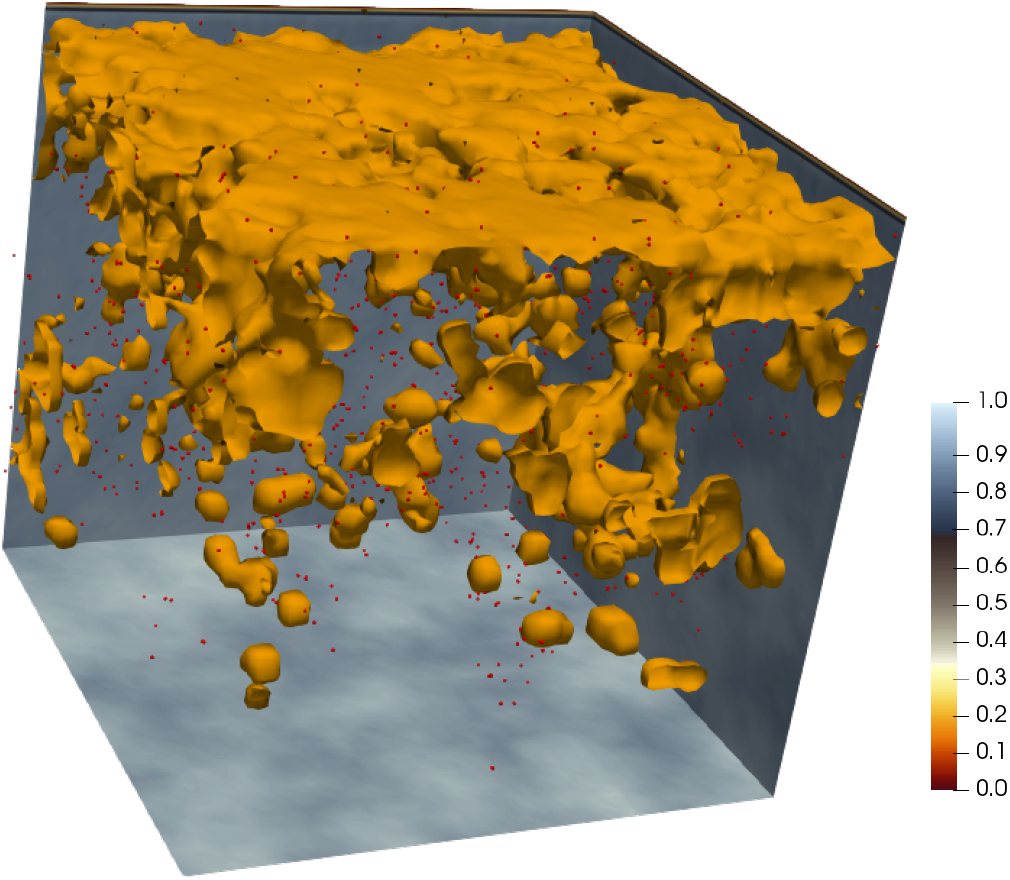
3D myoma invasion simulation. Shown is an indicative final time (t = 14 d) 3D model simulation result. The ECM density is shown in the background planes and corresponds to the colour bar on the right; the epithelial-like cancer cell density is visualised through contour surfaces corresponding to 0.1 of the average tumour density; the mesenchymal-like cancer cell particles are visualised as red dots. The experimental settings, initial conditions, and parameter values are discussed in Section 5 and Table 1. The size of the domain is 600 µm × 600 µm × 600 µm. A plethora of bio-medically relevant information can be extracted and studied from such simulation results, cf. Figures 8, 9 and Section 6.

**Figure 8:**
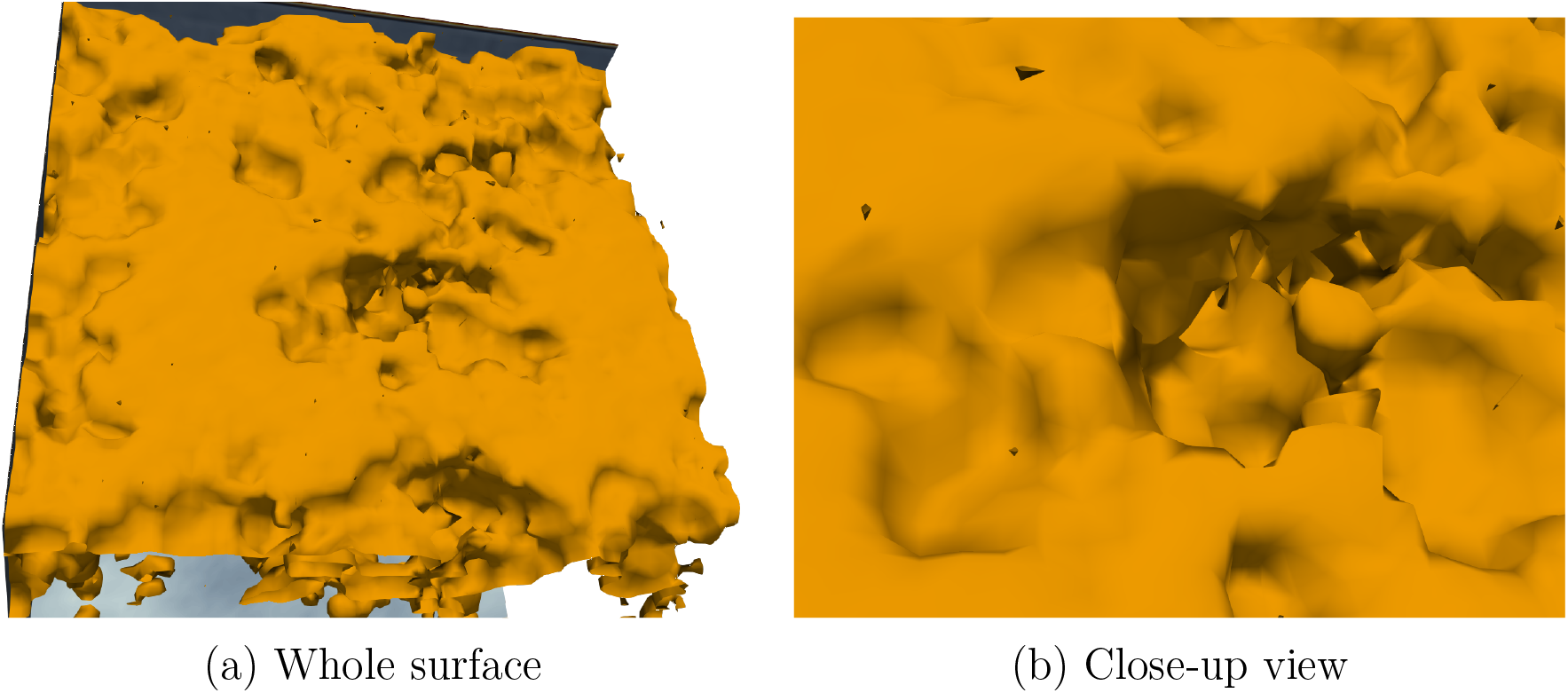
3D myoma invasion viewed from above. When viewed from above, the myoma invasion at day 14, which is also shown in Figure 7, presents the structure of the main body of the tumour as it grows due to proliferation and merging of the cancer cell “islands”. The detailed view (b) gives insight into the “inner” structure of the tumour. Like in previous figures, the epithelial-like cancer cell density is visualised through contour surfaces corresponding to 0.1 of the average tumour density.

**Figure 9:**
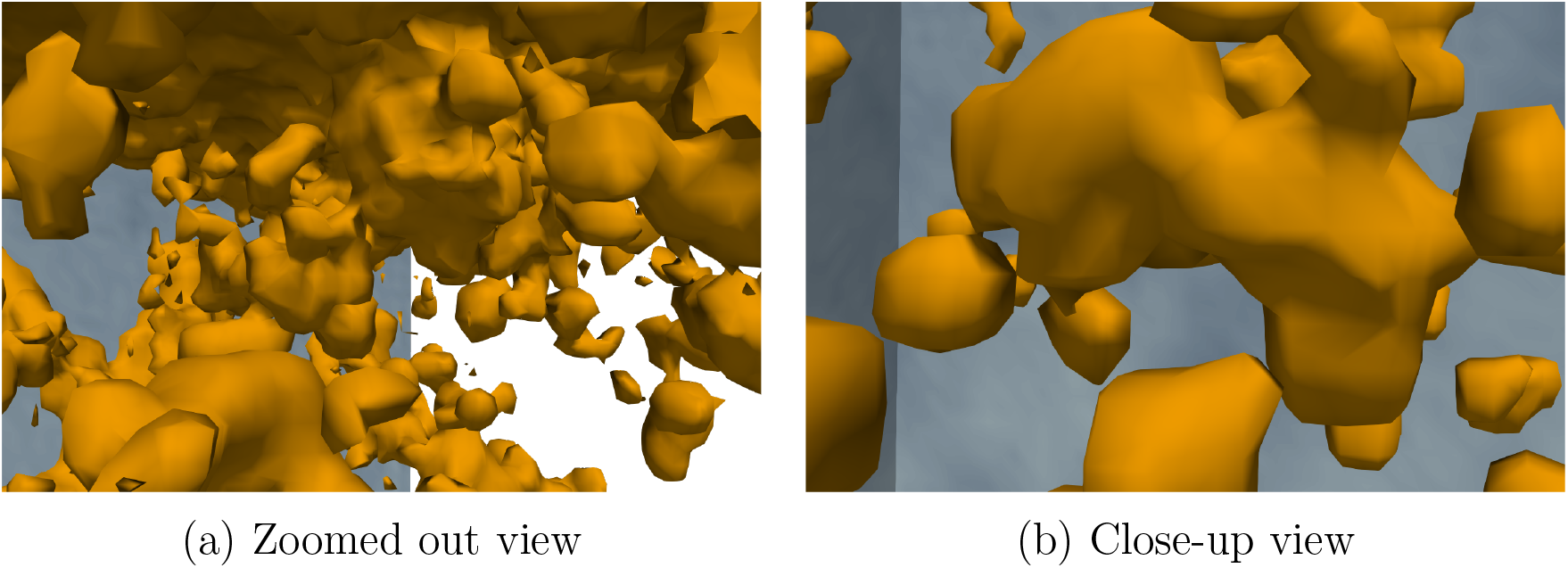
3D myoma invasion viewed from the inside. When viewed from inside the assay, the epithelial-like cancer cell density on day 14, which is also shown in Figure 7, reveals a variable structure of tumour growth. Newly formed epithelial-like invasion “islands” form and grow, and merge either with the initial/main body of the tumour (a) and/or with each other (b). Like in previous figures, the epithelial-like cancer cell density is visualised through contour surfaces corresponding to 0.1 of the average tumour density.

The migration of mesenchymal-like cancer cell particles takes the form of a haptotaxis-biased random motion, which is modelled by the system of SDEs (3.4). These cell particles escape from the main body of the tumour and, as they invade the ECM, they can undergo MET and give rise to new epithelial-like cell densities. The spatio-temporal evolution of these newly formed cell densities is then once again governed by the continuum model (3.1)–(3.3). Primarily due to proliferation and diffusion, these newly formed densities grow to become invasion “islands” that grow away from the non-invasive part of the tumour located on top of the ECM. According to the growth of these “islands”, they can be categorised into:

– Should they be sufficiently close to the top of the assay, these “islands” may merge with the upper layer of epithelial-like cancer cells on top of the ECM. They are then considered part of the non-invading tumour mass.
– The islands are considered as a part of the invading tumour if they grow at a sufficiently large distance from the top of assay so they have not merged with the non-invading tumour mass.

We refer also to Figure 4 for the description and definition of the invading versus non-invading cell area as applied both in the analysis of the experimental results by Nurmenniemi et al. (2009) as well as in our analysis of the model and simulation results.

### 5.1 Results

Through the simulations that we carried out, we found that the proposed three-dimensional hybrid model provides qualitatively and quantitatively biologically realistic results. This holds in particular, when the simulation results are compared against the organotypic *in vitro* HSC-3 invasion assays described in Nurmenniemi et al. (2009).

The simulation results shown in Figure 6 were conducted using the estimated parameter values in Table 1. Five of these, i.e. the parameters (4.1), were obtained by minimising the discrepancy between the simulations and the experimental measurements using a combination of the *maximal invasion depth*, the *total non-invading area*, and the *invasion index* “norms”. All of these are defined below.

In the experimental setting by Nurmenniemi et al. (2009), these quantities were measured as explained in Section 4.2 and Figures 3 and 4. The quantification of the experimental results after 14 days is visualised in Figure 2, yielding a median of 5.4700 × 10^−2^ µm for the maximal invasion depth of the epithelial-like cancer cells, and of 3.5270 × 10^−4^ µm^2^ and 3.6710 × 10^−4^ µm^2^ for their total non-invasive and invasive area, respectively.

Through the parameter estimation and sensitivity analysis processes, we have obtained the corresponding values 7.0130 × 10^−2^ µm, 3.9884 × 10^−4^ µm^2^ and 3.0398 × 10^−4^ µm^2^, respectively. It is worth noting that the maximal invasion depth in the experiments of Nurmenniemi et al. (2009) was measured as the mean of the three epithelial-like cancer cells that had invaded the domain the furthest. To ensure that our work represents the experimental measurements as closely as possible, we adopted this approach when measuring the simulation outcomes. To measure these particular quantities in our simulations, we first extracted nine vertical slices from the middle of the three-dimensional epithelial-like tumour density profiles. We then computed the above quantities in each of these slices as follows:

#### Non-invading area

We measure the non-invading area of the tumour as the area of the connected upper part of the epithelial-like density profile.

#### Invading area

The invading area of the tumour is computed by subtracting the non-invading area of the tumour from the overall area of epithelial-like density profile.

#### Maximum invasion depth

The maximum invasion depth is measured as the vertical distance between the invading epithelial-like cells and the lower boundary of the non-invasive area. To comply with the experimental methods, *cf*. Figure 3, we also compute the mean of the three largest invasion depths in every slice.

#### Invasion index

Like in Nurmenniemi et al. (2009), the invasion index is defined through the relation

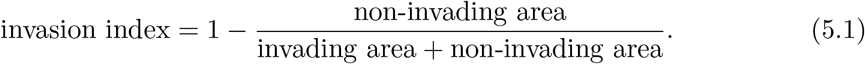

We use the above quantities, as well as (4.2), to compute the error/discrepancy between the experimental and simulation measurements. As these are absolute errors, they do not allow for a direct comparison between the corresponding quantities. So instead we define the relative errors

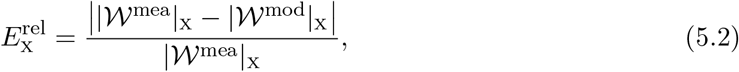

i.e. the ratio of the absolute error (4.2) between the experimental and simulation measurements over the experimental measurements. Representing by X = 1, 2, 3 the *maximum invasion depth*, the *invasion area*, and the *invasion index*, we determine the relative *root mean square* (RMS) of the errors 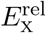 as

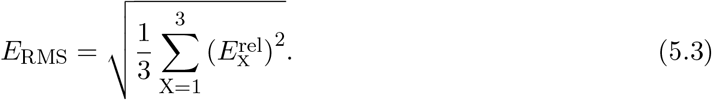

We used this relative RMS error (5.3) for the parameter estimation and sensitivity analysis of the model.

As an *a posteriori* study, we also estimated the *backwards and forwards sensitivity gradients*. These were obtained by varying each of the parameters to half (×0.5) and, respectively, double (×2) their reference value 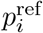, while retaining all other parameters at their reference values, and subsequently computing the sensitivity gradients (4.6). This way, we gained two additional pieces of information:

a. the signs of the backward and forward gradients indicate whether the reference value 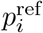 is (close to) the minimizer. A combination of negative-positive signs implies that the sensitivity decreases-increases around the reference value. A change in the sign of the sensitivity gradients implies that the sensitivity decreases and then increases, or vice versa, around the reference value.
b. The magnitudes of the two gradients indicate the sensitivity of the result to the variation of the parameter *p*_*i*_ along the corresponding direction.

#### Insights from the parameter sensitivity analysis

As discussed in Section 4.2.2, we slightly vary each of the model parameters 𝒫^sens^ (4.3)—one after the other—around their respective reference value 𝒫^ref^ (4.4). Due to the stochastic nature of the problem, the simulations were repeated 20 times for each parameter set. This is also highlighted in the Figures 10 and 11. The analysis yielded the following results, which are visualised in Figure 10:

**Figure 10:**
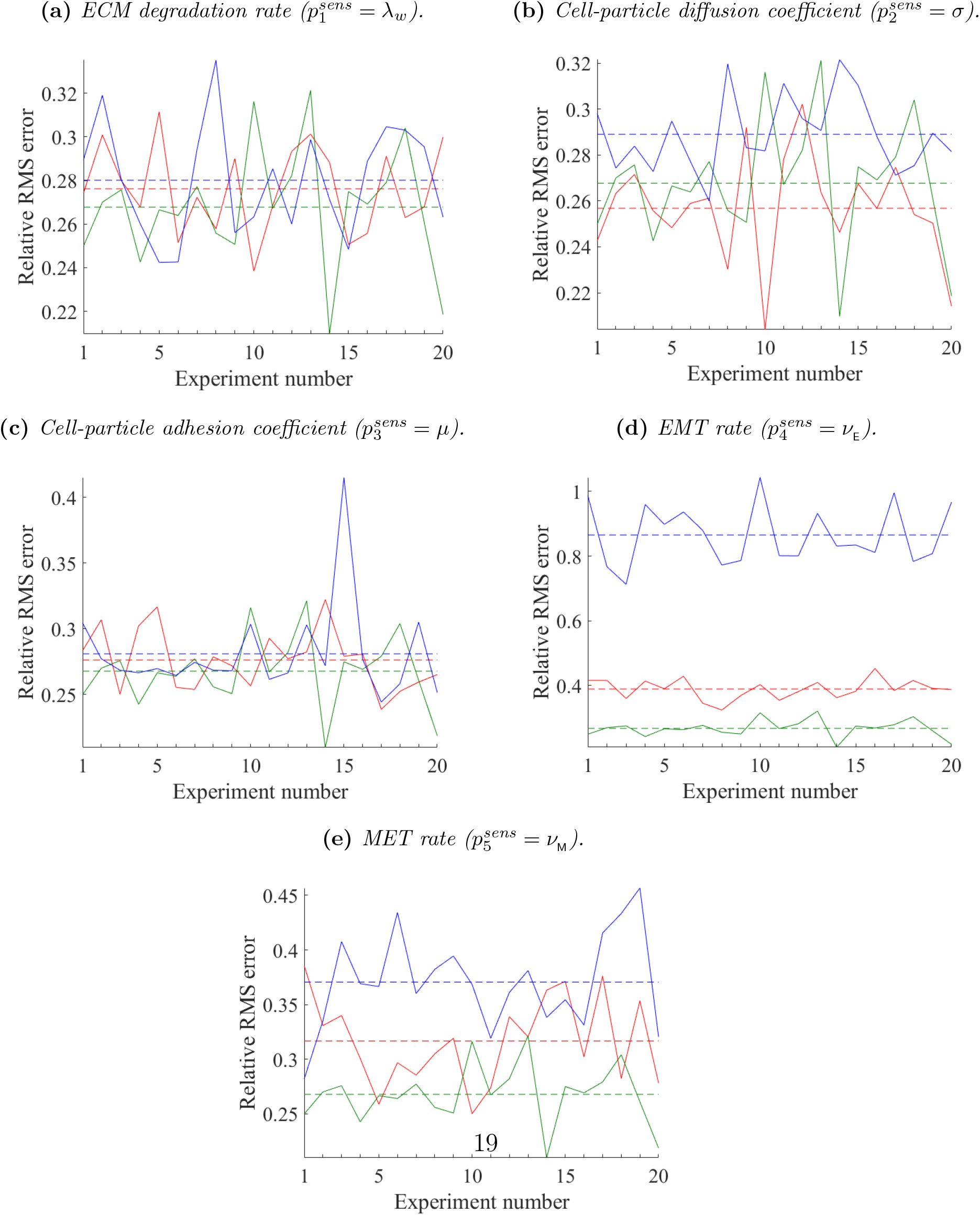
Sensitivity analysis. The subplots show the impact that the perturbation of each one of the five estimated parameters 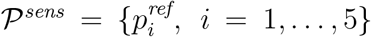 (4.3), from their reference values (green lines) to their perturbed states 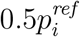 (red lines) and 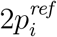 (blue lines) has on the relative RMS error (5.3) (vertical axes). Due to the inherent stochasticity of the model, each experiment is repeated 20 times (horizontal axes) for the same parameter set. The resulting average values are depicted by the corresponding dashed lines. In all subplots the reference states (green lines) are the same. It can be seen that the relative RMS error is more sensitive to variations of the EMT and MET rates shown in panels (d) and (e) than to the ECM degradation rates in panel (a). Similarly, we note that the average sensitivity of the relative RMS error to cell-particle diffusion and adhesion coefficients in panels (b) and (c) is similar.

ECM degradation rate 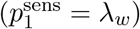: The sensitivity analysis results are shown in Figure 10a. The corresponding sensitivity gradients were -110.2882 and 80.5483, respectively, for the experiments conducted with values 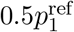 and 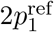 with 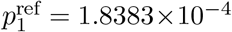, M cm^−3^d^−1^. The change in the sign of the sensitivity gradients implies that the minimum of the relative RMS error (5.3) is attained around the reference value 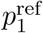. This is an indication that the reference parameter was sufficiently well estimated. Furthermore, the strong gradients imply that the relative RMS error is quite sensitive to the ECM degradation rate *λ*_*w*_.

Cell particle diffusion coefficient 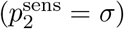: The sensitivity analysis results are shown in Figure 10b. The sensitivity gradients were 0.0033 and 0.0032, respectively, for the experiments conducted with values 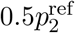 and 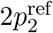 with 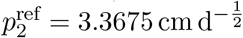. The two positive signs imply that the error increases with the parameter. This should serve as an indication to reduce the reference parameter value 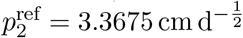 and to perform the sensitivity analysis again thereafter. However, as the magnitude of the gradients—and hence the sensitivity to this particular parameter—is small, no strong benefit is to be expected. Hence, we conclude that the reference parameter value 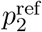 is acceptable.

Cell particle drift coefficient 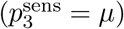: The sensitivity analysis results are shown in Figure 10c. The corresponding sensitivity gradients were -0.3658 and 0.2830 for the experiments conducted with values 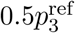 and 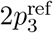 with 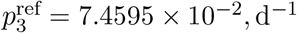. As in the case of the ECM degradation rate *λ*_*w*_, the change in sign indicates that the minimiser is (close to) the reference value 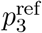. However, by direct comparison to the ECM degradation rate *λ*_*w*_, we deduce that the relative RMS error (5.3) is less sensitive to this parameter.

EMT rate 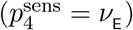: The sensitivity analysis results are shown in Figure 10d. The sensitivity gradients were -3.3807 and 8.2855, respectively, for the experiments conducted with values 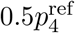 and 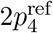 with 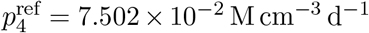. Again, the change in sign indicates that the minimiser of the relative RMS error is attained around the reference value 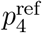. Moreover, the relative RMS error (5.3) is more sensitive to the EMT rate *ν*_E_ than to the particle drift coefficient *σ* but less sensitive than to the ECM degradation rate *λ*_*w*_.

MET rate 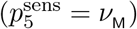: The sensitivity analysis results are shown in Figure 10e. The sensitivity gradients were -0.2091 and 0.2201, respectively, for the experiments conducted with values 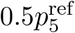 and 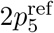 for 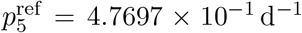. As before, the change in sign indicates that the minimiser of the relative RMS error is (close to) the reference value 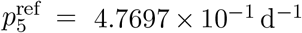. Moreover, the relative RMS error is less sensitive to the MET rate *ν*_M_ than to the ECM degradation rate *λ*_*w*_, the particle drift coefficient *µ*, and to the EMT rate *ν*_E_.

As summarised in Table 2, the sensitivity analysis has shown that the parameter values *P* ^sens^ (4.3) that we determined through the parameter estimation, *cf*. Table 1, are sufficiently close to the minimiser of the relative RMS error. Hence, they are a good fit to the model. Furthermore, the relative RMS error shows different degrees of sensitivities to the different parameters. Ordering the five parameters according to the apparent sensitivity of the relative RMS error, we get (from least to most): particle diffusion coefficient *σ*, MET rate *ν*_E_, particle adhesion coefficient *µ*, EMT rate *ν*_E_, and ECM degradation rate *λ*_*w*_.

**Table 2:** Summary of the backward and forward sensitivity gradients with respect to the five model parameters 𝒫^sens^ (4.3) around their respective reference value 𝒫^ref^ (4.4). The ± sign of the gradients indicates the direction of increase or decrease respectively of the relative RMS error (5.3), and their magnitude the absolute sensitivity of the relative RMS error to changes of the corresponding parameter.

|  | backward | forward |
| --- | --- | --- |
| $\lambda_w$ | -110.2882 | 80.5483 |
| $\sigma$ | 0.0033 | 0.0032 |
| $\mu$ | -0.3658 | 0.2830 |
| $\nu_E$ | -3.3807 | 8.2855 |
| $\nu_M$ | -0.2091 | 0.2201 |

It is worth noting that we have opted to minimise the relative RMS error (5.3), rather than its constituent relative errors *e*_*i*_, *i* = 1, 2, 3, as we aimed to account for all of the quantitative output provided in Nurmenniemi et al. (2009). Still, had we performed the sensitivity analysis against one of these three quantities separately, we would have extracted information that describes how this particular quantity depends on the parameters under discussion. In particular, Figure 11 exhibits the dependence of the maximum invasion depth of the epithelial-like cancer cells on the cell particle diffusion coefficient *σ*. After performing 20 simulation experiments for each parameter set, it can be clearly be seen that increasing *σ* from the reference value 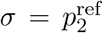 (green line) to 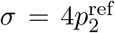 (blue line) causes the average maximum invasion depth to increase only slightly. On the contrary, when *σ* decreases from the reference value 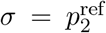 (green line) to 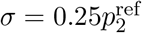 (red line) the average maximum invasion depth decreases significantly. As a result, the average maximal invasion depth is closer to the median value 5.4700 × 10^−2^ µm measured experimentally by Nurmenniemi et al. (2009), and hence reducing 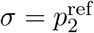 might lead to a better fit to the experimental data. Yet, comparing with the results from the sensitivity analysis that accounted for the relative RMS error rather than solely for the maximum invasion depth, which is shown in Figure 10b, we find that decreasing the particle diffusion coefficient does not benefit significantly in decreasing the relative RMS error.

## 6 Perspectives

We have proposed a three-dimensional model based on the two-dimensional model introduced in Sfakianakis et al. (2020), that accounts for the phenotypic variation of cancer cells by distinguishing between an epithelial-like and a mesenchymal-like phenotype. The model addresses dynamic mutations between these two cell phenotypes in the form of EMT and its reverse process MET.

The newly developed model is a three-dimensional, genuinely hybrid atomistic-continuum combination of macroscopic densities and microscopic atomistic profiles. The former represents the epithelial-like part of the tumour, the ECM, and the MT1-MMPs, which obey a system of PDEs. The latter represent isolated mesenchymal-like cancer cells whose time evolution is governed by a system of SDEs. The coupling between the two cellular phenotypes takes the form of a phase transition between continuum and discrete quantities.

**Figure 11:**
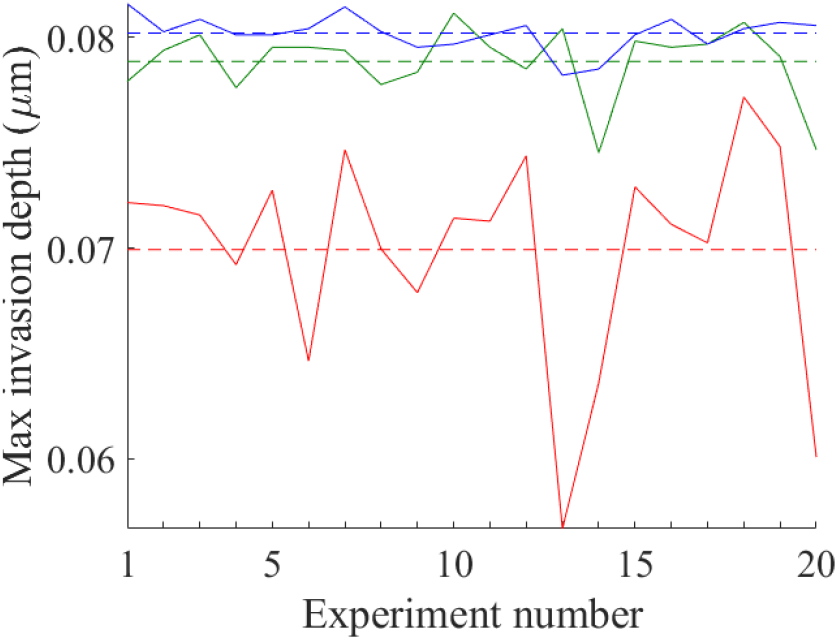
Sensitivity analysis for particle diffusion coefficient against maximum invasion depth of the epithelial-like cancer cells. For the cell particle diffusion coefficient 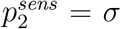, the maximum invasion depth of the epithelial-like cancer cells was computed for 20 simulations of the experiment. In the plot, the results obtained from the reference parameter set 𝒫^ref^ are shown in green; the results from parameter sets with the parameter values 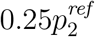 and 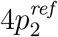 are shown in red and blue, respectively. The means were computed and represented by a horizontal dashed line in the respective colour. For result interpretation, see text.

The model is primarily informed by a number of parameters extracted from the relevant biological literature. Still, the five model parameters (4.3) cannot be inferred directly from the literature. Instead, they are estimated by a combination of global and local inverse parameter estimation methods augmented by an *a posteriori* local parameter sensitivity analysis. To this end, the model is tested against an *in vitro* organotypic invasion assay experiment by Nurmenniemi et al. (2009), where the invasion of OSCC cells into uterine leiomyoma tissue was studied.

We find the resulting model predictions to be in good qualitative and quantitative agreement with the experimental findings. This allows us to draw conclusions with respect to the impact of several of the model components. In particular, we found that the ECM degradation rate *λ*_*w*_ is the most influential among these five parameters, followed by the EMT rate *ν*_E_. We also observed that increasing either the particle diffusion coefficient *σ* or the EMT rate *ν*_E_ leads to an increase of the invasion through increased numbers of invasion “islands” and an increase in the volume of the main tumour body, respectively, *cf*. Figure 12.

**Figure 12:**
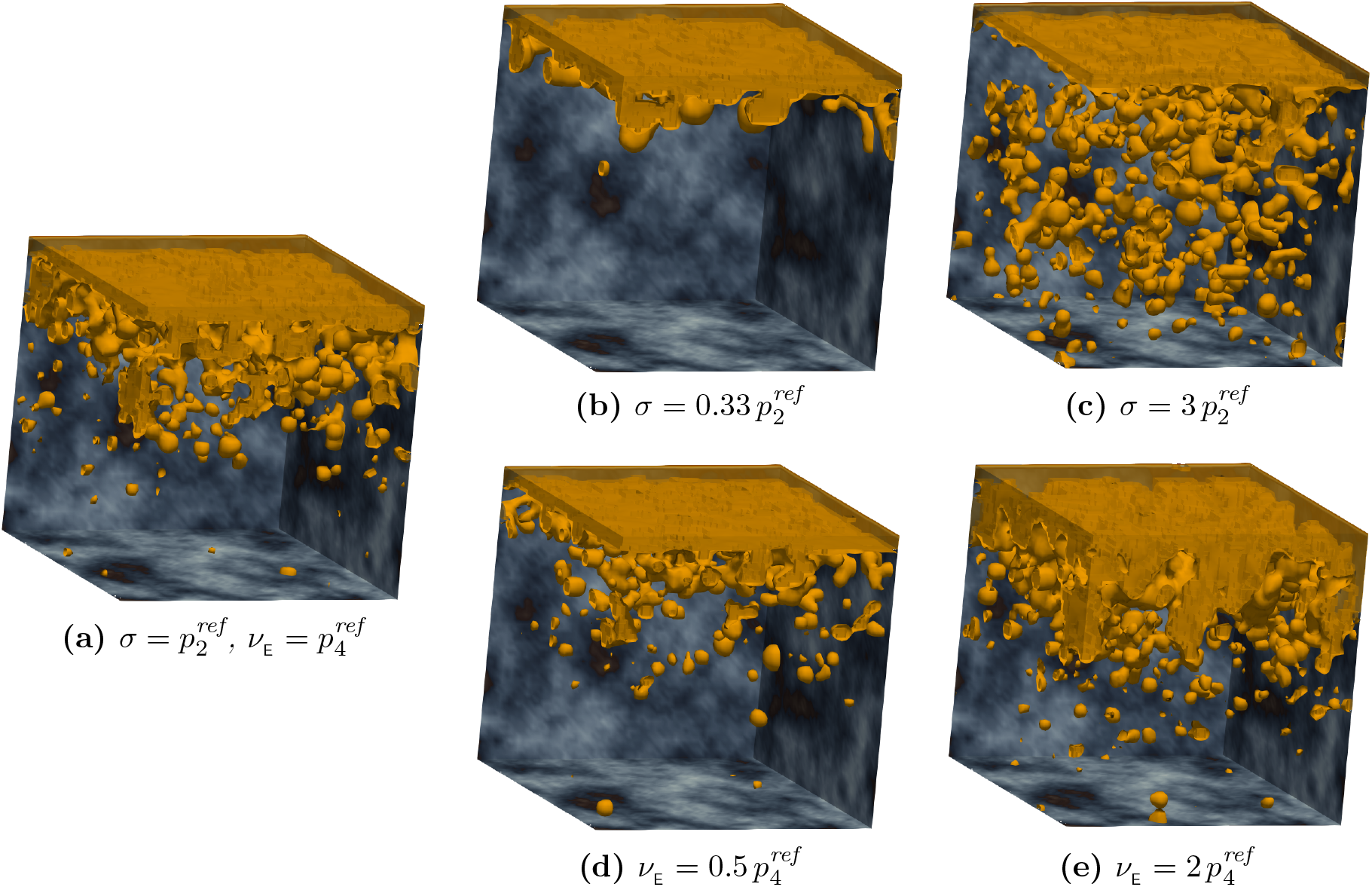
The impact of the cell particle diffusion σ and the EMT rate ν_E_. **(a)**: Simulation using the reference parameter set, cf. Table 1, and in particular 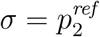and 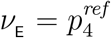. **(b)** and **(c)**: We varied σ from the reference value to 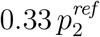 and 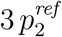, respectively, while maintaining the rest of the parameters as in Table 1. These simulations exhibit that the invasion increases with σ, primarily through increased numbers of invasion “islands” away from the main body of the tumour. **(d)** and **(e)**: We varied ν_E_ from the reference value to 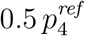 and 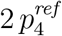, respectively, while retaining the other parameters as in Table 1. These simulations show increased invasion as ν_E_ increases. However, as opposed to **(b)** and **(c)**, this stems primarily from the increase of the volume of the tumour’s main body.

In this paper we focussed on the invasion of OSCC, the most common type of *head and neck squamous cell carcinoma* (HNSCC) (Kim et al., 2019). In HNSCC, distal organ metastasis occurs comparatively rarely compared to other cancers. Instead, local progression is a major cause of HNSCC-related mortality (Chang et al., 2013). A study by Chang et al. (2013) suggests that the mechanism behind this is the induction of MET and inhibition of migration in HNSCC cells—and, amongst others, HSC-3 cells in particular—through connective tissue growth factors in the microenvironment of a primary tumour. Correspondingly, an often-observed phenomenon in OSCC, as well as other types of carcinomas (Japanese Gastric Cancer Association, 2011; Ito et al., 2012; Masuda et al., 2017), is the occurrence of “islands” of cancer cells outside of the main body of the tumour (Nurmenniemi et al., 2009; Almangush et al., 2018). In other types of carcinomas, the involvement of MET in dissemination and metastatic colonisation is of crucial importance (Dongre and Weinberg, 2019). Therefore, to apply the model to other types of carcinomas, we will account for the metastatic spread from one site of the body to other sites through multiple domains in a multi-organ model, *cf*. Franssen et al. (2019) and Franssen and Chaplain (2019). For each organ with primary and secondary spread, the parameter settings can be adjusted according to its microenvironment.

Also, to make the movement of the single mesenchymal-like more biologically realistic, a further extension of this modelling framework is the improved description of the migration of mesenchymal-like cancer cells. A lamellipodium-based cell-migration model such as the one presented in Sfakianakis et al. (2018), could replace the ad-hoc SDEs that we use this work. The various forms of cell-cell interactions, which in the current model have been reduced to the mere minimum of competing for free space/resources, are another component to account for in the model in the future. Moreover, we aim to incorporate diffusible MDEs such as MMP-2, rather than relying solely on the action of the membrane-bound MT1-MMP for the degradation of the ECM.

## Appendices

### A Phase transition operators between densities and particles

#### Particle-to-density transition operator for MET

Let 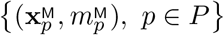 be a collection of particles that represent mesenchymal-like cancer cells. Using (3.6) and (3.7), we define the *particle-to-density* operator ℱ as

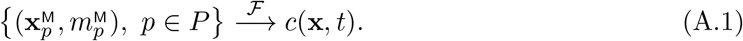

To define the function *c*(**x**, *t*), we go through all the particles that represent mesenchymal-like cancer cells and consider their corresponding density formulation according to (3.5). The support *K*_*p*_ of every particle overlaps with several of the partition cells *M*_*i*_, *i* ∈ *I*. We assign the corresponding portion of the particle mass to every partition cell *M*_*i*_:

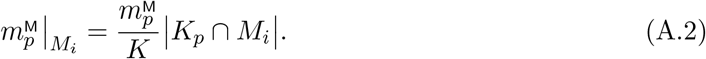

In a similar fashion, we account for the contribution of all particles *p* ∈ *P* to a partition cell *M*_*i*_:

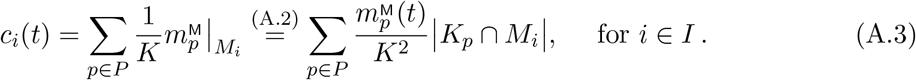

In view of equations (3.5) and (A.3), we deduce the density function *c*(**x**, *t*) over the full domain Ω to be

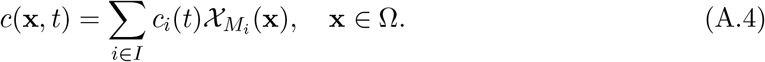

For simplicity, we assume that the mesenchymal-like cancer cell particles 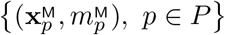 undergo MET to become epithelial-like cancer cells (below abbreviated as *ECC*) randomly through the process

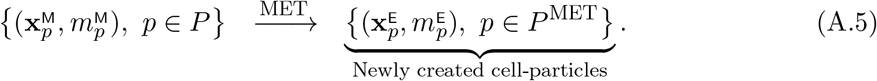

The resulting epithelial-like cancer cell particles are instantaneously transformed to density via the *particle-to-density* operator ℱ given in equation (A.1):

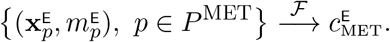

Consequently, the MET can be expressed in operator form as

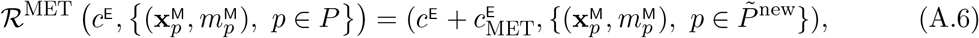

where 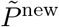 is a re-enumeration of the set difference *P \ P* ^MET^.

#### Density-to-particle transition operator for EMT

Given a density function *c* = *c*(**x**, *t*), we define the *density-to-particle* operator *B* for a general particle as

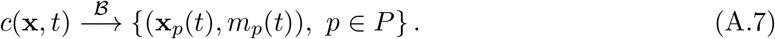

We assign one particle with mass

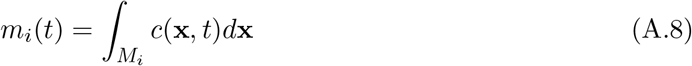

and position

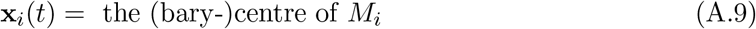

to every cube partition cell *M*_*i*_, *i* ∈ *I*.

The density-to-particle transition is used in this paper to model the EMT process. As in the case of MET in Section A, EMT is—at this stage—represented using a simplified approach where a randomly chosen part of the epithelial-like cancer cells (in density formulation) 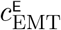 undergoes EMT to give rise to mesenchymal-like cancer cells. For this, the full domain is discretised into partition cuboids *M*_*i*_, *i* ∈ *I*. EMT takes place with some probability in each cuboid that contains some material. The larger the amount of material in a cuboid, the higher the probability that one cell undergoes EMT. We perform this process in steps. First, the randomly chosen part of the epithelial-like cancer cell density 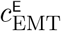 transitions to mesenchymal-like cancer cell density:

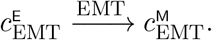

This mesenchymal-like cancer cell density is immediately transformed to mesenchymal-like cancer cell-particles via the *density-to-particle* operator ℬ given in equation (A.7):

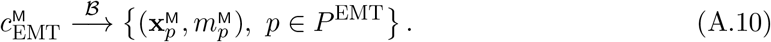

Here 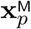 and 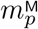 are given by equations (A.8) and (A.9), respectively, and *P* ^EMT^ is the set of indices corresponding to the particles that perform EMT. Subsequently, the family of existing mesenchymal-like cancer cell particles—below abbreviated as *MCC* —is updated with these newly created particles. It is hence given by the disjoint union

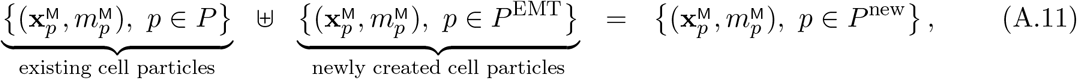

where *P* ^new^ is a re-enumeration of the multiset 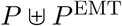.

Overall, the EMT operator consequently reads as

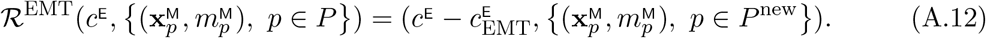

